# Particle-Swarm Based Modelling Reveals Two Distinct Classes of CRH^PVN^ Neurons

**DOI:** 10.1101/2022.03.22.485381

**Authors:** Ewandson L. Lameu, Neilen P. Rasiah, Dinara V. Baimoukhametova, Spencer Loewen, Jaideep S. Bains, Wilten Nicola

## Abstract

Electrophysiological recordings can provide detailed information of single neurons’ dynamical features and shed light into their response to stimuli. Unfortunately, rapidly modeling electrophysiological data for inferring network-level behaviours remains challenging. Here, we investigate how modeled single neuron dynamics lead to network-level responses in the paraventricular nucleus of the hypothalamus (PVN), a critical nucleus for the mammalian stress response. Recordings of corticotropin-releasing hormone neurons from the PVN (CRH^PVN^) were performed using whole-cell current-clamp. These, neurons, which initiate the endocrine response to stress, were rapidly and automatically fit to a modified Adaptive Exponential Integrate and Fire model (AdEx) with Particle Swarm Optimization (PSO). All CRH^PVN^ neurons were accurately fit by the AdEx model with PSO. Multiple sets of parameters were found that reliably reproduced current-clamp traces for any single neuron. Despite multiple solutions, the dynamical features of the models such as the rheobase current levels, fixed points, and bifurcations, were shown to be stable across fits. We found that CRH^PVN^ neurons can be divided into two sub-types according to their bifurcation at the onset of firing: saddles (integrators) and sub-critical Hopf (resonators). We constructed networks of these fit CRH^PVN^ model neurons to investigate the network level responses of CRH^PVN^ neurons. We found that CRH^PVN^-resonators maintain baseline firing in networks even when all inputs are inhibitory. The dynamics of a small subset of CRH^PVN^ neurons may be critical to maintaining a baseline firing tone in the PVN.

**Key Points:**

- Corticotropin-releasing hormone neurons (CRH^PVN^) in the paraventricular nucleus of the hypothalamus act as the final neural controllers of the stress response.
- We developed a rapid computational modeling platform that uses Particle-Swarm Optimization to rapidly and accurately fit biophysical neuron models.
- A model was fit to each patched neuron without the use of dynamic clamping, or other procedures requiring sophisticated inputs and fitting procedures. Any neuron undergoing standard current clamping for a few minutes can be fit by this procedure
- The dynamical analysis of the modeled neurons shows thatCRH^PVN^ comes in two specific ‘flavours’: CRH^PVN^-resonators and CRH^PVN^-integrators.
- Network simulations show thatCRH^PVN^-resonators are critical to retaining the baseline firing rate of the entire network of CRH^PVN^ neurons as these cells can fire rebound spikes and bursts in the presence of strong inhibitory synaptic input.

## 1 Introduction

When faced with an impending stressor, or threat, the Hypothalamic-Pituitary-Adrenal (HPA) axis of the neuroendocrine system coordinates an immediate response to prepare an organism for potential action [1]. The final neural controllers of the classical stress response in the hypothalamus are the corticotropin releasing hormone neurons of the paraventricular nucleus of the hypothalamus (CRH^PVN^). These neurons increase their activity in response to stress, which causes the release of CRH into the median eminence where it eventually stimulates the pituitary gland. This cascade ultimately results in glucocorticoid production from the adrenal glands and comprises the hormonal response to stress.

CRH^PVN^ neurons are situated atop the hierarchy of this complex neuroendocrine chain. Their position as the master controllers make these neurons uniquely exploitable by upstream nuclei to initiate, attenuate, block, or otherwise modify the body’s response to stress. Indeed, these neurons are heavily innervated by GABAergic inputs as a potential source of suppression in the stress response [2]. Direct activation or deactivation of CRH^PVN^ neurons with optogenetic stimulation can change the behavioural profile of animals [3, 4], indicating that these neurons may be directly relevant for coordinating behavioural responses to stress. Finally, stress-related pathologies, such as hypertension or generalized anxiety disorder have been strongly linked to these neurons [5, 6].

Although CRH^PVN^ neurons are the final neural-controllers of the HPA-axis, the dynamics of isolated neurons, and the potential behaviours of these neurons in a network have yet to be directly modeled. This is for two primary reasons. First, CRH^PVN^ neurons do not form a coherent nucleus of cells within the PVN and are diffusively distributed among other non-CRH^PVN^ neurons. This makes simultaneously patching many neurons for a large scale analysis difficult, and multi-electrode recordings which would randomly sample from all PVN neurons, non selective. Second, modelling neurons with current clamp protocols is often a lengthy process, requiring dynamic clamping or long and noisy current injections to adequately estimate the voltage-current relationships in neural dynamics [7].

To unravel the mechanisms behind the mammalian response to stress, we performed whole cell recordings of identified CRH^PVN^ neurons using two different current clamp protocols. The neurons were recorded in brain slices from mice with different social interactions, housing characteristics, acute and chronic stress exposures, exogenous glucocorticoid exposure, and sex. The patched neurons were fit using a modified Adaptive Exponential integrate-and-fire neuron model (AdEx) [8, 9] with particle swarm optimization [10–14] used to determine the electrophysiological parameters of the AdEx model. As all the modifications to the AdEx model used here were to the spiking characteristics, all prior bifurcation analyses of the classical AdEx model were directly applicable. We found two distinct classes of CRH^PVN^ neuron: CRH^PVN^-resonators and CRH^PVN^-integrators. The resonators displayed an ultra-slow (<0.5 Hz) resonance frequency, and were found to be critical to maintaining the baseline firing response. Further, we found that female mice undergoing a chronic stress protocol increased their proportion of resonators, where as males did not. Network simulations show that CRH^PVN^-resonators can maintain firing despite an overwhelming amount of simulated inhibitory input, pointing to their potential role as a critical network component maintaining baseline firing in stress circuits. Our results show that CRH^PVN^ neurons are a dynamic population of cells that evolve in response to the stressors, and display a diversity of dynamical behaviours.

## 2 Materials and Methods

The experimental protocols used in the current-clamp recordings of CRH neurons are described in section 2.1 while the Adaptive Exponential (AdEx) integrate-and-fire model and the modifications we propose to improve the reproduction of spike features observed in experiments are described in section 2.2. Next, the Particle-Swarm-Optimization (PSO) parameter search is explained together with the particles’ movement equations and the parameter bounds used to restrict the search in section 2.4. The specific cost function used in PSO, which incorporates both spike times and voltage traces for parameter fitting is described in 2.5. Finally, we describe the analytical equations used to extract the dynamical features of CRH^PVN^ neurons from the fit models in 2.6.

### 2.1 Current Clamp Recordings of Corticotropin-Releasing-Hormone Neurons

The electrophysiological data consists of *in vitro* current clamp recordings of Corticotropin-Releasing Hormone (CRH^PVN^) neurons from the Paraventricular Nucleus of Hypothalamus (Figure 1A). During a threat or stressful situation, CRH^PVN^ neurons increase their activity and release CRH^PVN^. This eventually promotes the multifaceted mammalian response to stress [21, 22].

Mice were anesthetized with isoflurane and decapitated. The brain was rapidly extracted and submerged in freezing slicing solution (0°C, 95% *O*_2_, 5% *CO*_2_ saturated) containing 87 mM *NaCl*, 2.5 mM *KCl*, 0.5 mM *CaCl*_2_, 7 mM *MgCl*_2_, 25 mM *NaHCO*_3_, 25 mM D-glucose, 1.25 mM *NaH*_2_*PO*_4_ and 75 mM sucrose. Coronal slices were taken at 250μm thickness using a vibratome (Leica). Slices containing the PVN were hemisected and were incubated for 1 hour in artificial cerebral spinal fluid (aCSF) (30°C, 95% *O*_2_, 5% *CO*_2_ saturated) containing 126 mM *NaCl*, 2.5 mM *KCl*, 26 mM *NaHCO*_3_, 2.5 mM *CaCl*_2_, 1.5 mM *MgCl*_2_, 1.25 mM *NaH*_2_*PO*_4_ and 10 mM glucose. They were then transferred to a recording chamber super-fused with aCSF (95% *O*_2_, 5% *CO*_2_ saturated) at a constant rate of [1, 2] ml *min*^-1^. The temperature within the chamber was held constant at [30°, 32°]C. Slices containing the PVN were visualized using an Olympus BX51WI upright microscope fitted with infrared differential interference contrast optics. Patch pipettes were pulled from borosilicate glass pipettes ([3, 6] MΩ tip resistance), and were filled with internal solution containing 108 mM potassium gluconate, 2 mM *MgCl*_2_, 8 mM sodium gluconate, 8 mM *KCl*, 1 mM *K*_2_ — *EGTA*, 4 mM *K*_2_ — *ATP*, 0.3 mM *Na*_3_ — *GTP* and 10 mM HEPES. CRH^PVN^ neurons from CRH^PVN^-CreTdTomato mice were identified by expression of TdTomato fluorescent marker (previously characterized by [23]). Signals were amplified (Multiclamp 700A, Molecular Devices), low pass filtered at 1 kHz, digitized at 100 kHz (Digidata 1322, Molecular Devices), and recorded (pClamp 9.2, Molecular Devices) for offline analysis.

The current was clamped through different levels for each CRH^PVN^ neuron. In total, 248 CRH^PVN^ neurons were recorded and divided into 9 samples representing different stressor conditions and animal sex, under 2 current clamp protocols:

**Protocol 1:** Female Footshock (27 neurons), Male Footshock (22 neurons), Female Group Housed (47 neurons), and Female Single Housed (55 neurons). Cells were held at initial holding voltage of –70mV. Each sweep lasted 3s and contained an initial hyper-polarizing pre-pulse (–30pA, 0.5s) immediately followed by incremental depolarizing steps of 20pA (range [0, 180] pA, 0.5s).
**Protocol 2:** Male no-stress (26 neurons), Female Control (17 neurons), Female CORT (15 neurons), Male Control (19), and Male CORT (20 neurons). Cells were held at initial holding voltage of −70mV, and received 0.5s currents steps of 10pA (range [−40, +70]pA) at 1Hz. Regarding the animals under CORT Delivery, CRH-CretdTomato and CRH-Cre mice (p30, p50), were used for electrophysiological experiments. Animals were singly housed in a colony room with 12:12 hour light-dark cycle. Mice were given ad libitum access to a 0.95% *EtOH* solution containing 25*μ*g/ml CORT as their sole source of drinking water for 7 days. To prepare the solution, corticosterone (Sigma-Aldrich), was first dissolved in 95% ethanol and sonicated for 1 – 2min before dilution with tap water to 25*μ*g/ml. Solutions were replaced every 2 – 3 days and the amount of solution consumed was recorded. All water bottles containing solution were wrapped with foil to prevent light exposure.

### 2.2 The Modified Adaptive Exponential Integrate-and-Fire Model

The current clamp recordings of CRH^PVN^ neurons will form a basis with which to fit a straight-forward modification of the Adaptive Exponential Integrate and Fire Model (AdEx) proposed by [8, 9]. We note that similar modifications have been used by other groups (e.g. [24–26]). These modifications more faithfully reproduce neural dynamics from electrophysiological data, such as variable spike-thresholds or spike amplitudes.

In total, we added three new equations (Equation 3), similarly to those proposed by [26]. These new terms enable the model to dynamically evolve spike features as a result of prior spiking history, which was observed in data. In particular, the spike threshold *V_T_*, spike amplitude *V_p_* and reset voltage *V_r_* were observed to dynamically vary with successive spikes. The five modified-AdEx model equations are given by:

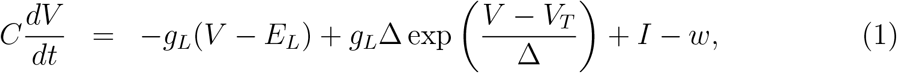

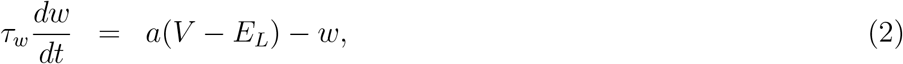

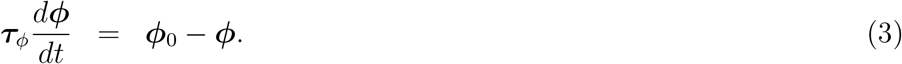

Where *ϕ* = [*V_T_*, *V_r_*, *V_p_*] and *ϕ*_0_ their respective resting values [*V*_*T*_0__, *V*_*r*_0__, *V*_*p*_0__]. The parameters listed are the membrane capacitance *C*, the external or applied current

**Figure 1:**
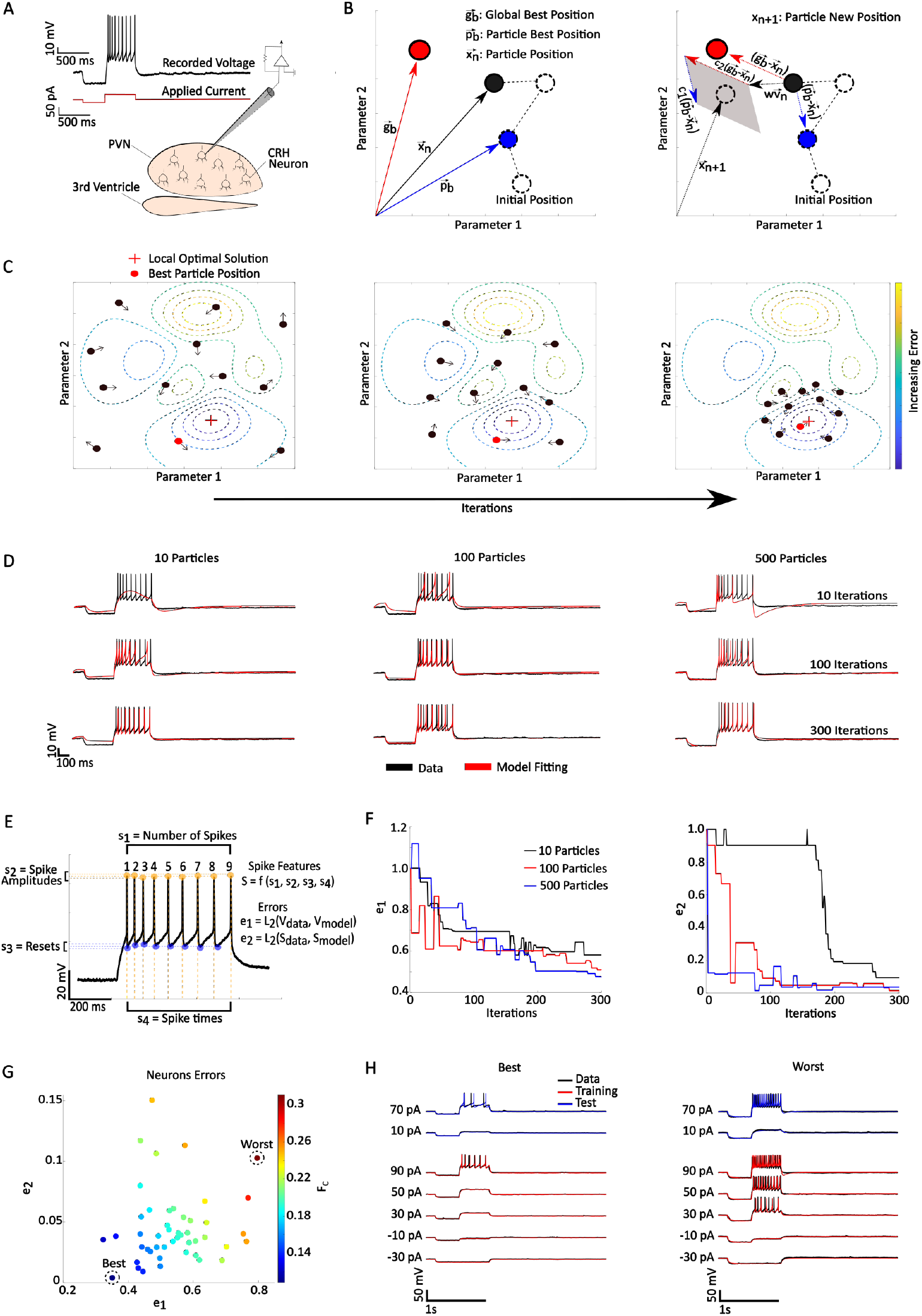
Using particle swarm optimization to fit models to CRH^PVN^ neurons. **(A)** schematic illustration of a PVN slice preparation and CRH^PVN^ neurons Current clamp record from a single neuron (black) receiving an external current (red). **(B)** On the left a PSO illustration of a particle moving in parameter space, *x_n_* represents its actual position, *p_b_* its best position (smallest *F_c_* achieved) and *g_b_* the best position among the swarm. The gray area highlights the possible new position *x*_*n*+1_ according to the movement equations of the particle. **(C)** Swarm search in the parameter space, the red cross depicts the global minima and the red particle is the best position overall. The arrows shows the movement direction. **(D)** Fitting variations according to the number of particles and iterations, the black lines represent the real data and in red the model. **(E)** Error function elements, *e*_1_ is the error associated with the voltage trace differences, and *e*_2_ is the error associated with spike features such as *s*_1_ number of spikes, *s*_2_ spike amplitudes, *s*_3_ reset voltage and *s*_4_ spike times. **(F)** Iteration evolution of errors *e*_1_ (left) and *e*_2_ (right) for 10 (black), 100 (red) and 500 (blue) particles. **(G)** Pair of errors of each neuron fit from FG sample, the dashed circles highlight the neuron with best (smallest *F_c_*) and the worst fit (highest *F_c_*) of the sample. **(H)** Data neuronal traces (black) of the best (left) and worst(right) fit neuron in the FG sample, in red the traces used by the PSO algorithm to find the model parameters (training) and in blue the testing traces.

*I*, the leak reversal potential *E_L_* and leak conductance *g_L_*, an exponential slope factor Δ (which controls spike width), an adaptation time constant *τ_w_*, and the level of sub-threshold adaptation *a*. The latter two parameters control the cells ability to reduce the rate of subsequent spikes as a function of spike history (e.g. spike frequency adaptation). Since *V_T_*, *V_r_* and *V_p_* are now variables, *τ_ϕ_* stands for their respective time constants *τ_ϕ_* = [*τ_T_*, *τ_r_*, *τ_p_*].

When the membrane potential of neuron is above a threshold *V* ≥ *V_p_*, the state variables are updated according to the following the rules:

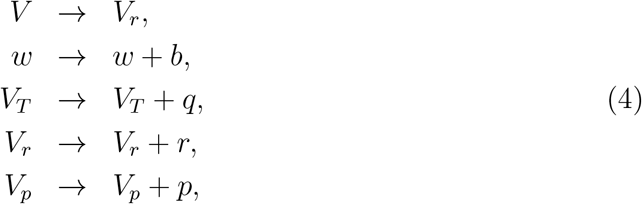

The external current, *I*, was provided by the current clamp parameters under the two separate protocols. Thus, the model consists of a set of *M* = 16 parameters to be fit given by the parameter vector ***x*** = [*C*, *g_L_*, *E_L_*, Δ, *τ_w_*, *a*, *τ_T_*, *τ_r_*, *τ_p_*, *b*, *q*, *r*, *p*, *V*_*T*0_, *V*_*r*0_, *V*_*p*0_]. The models, fitting procedure, and data analysis, were performed in Matlab2020b [27] using a standard Euler Method with 0.05 ms integration step. An example of the dynamics of each AdEx variable for a single neuron fit is depicted in Figure 2.

**Figure 2:**
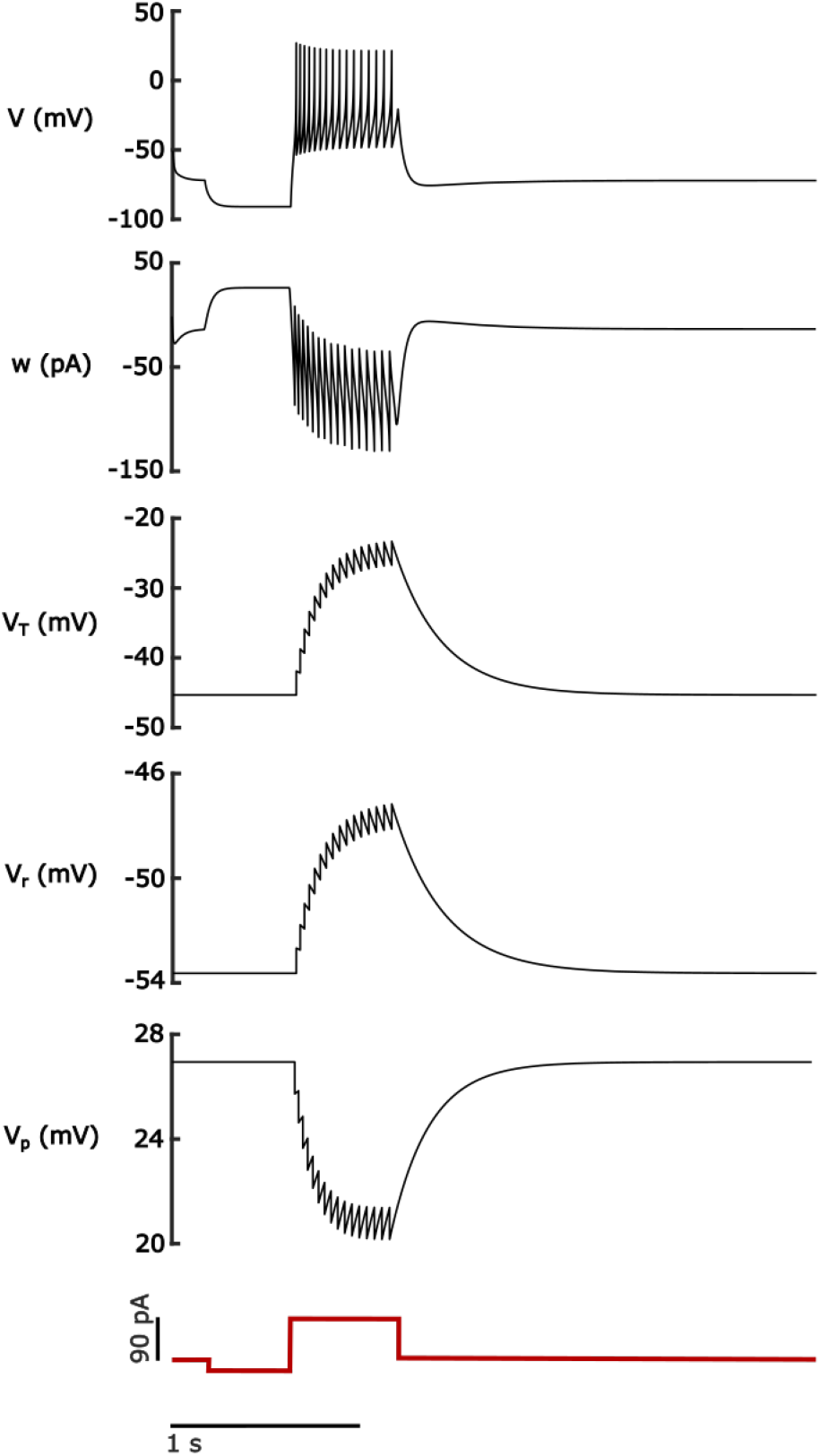
Modified adaptive exponential integrate-and-fire model parameter evolution. Time evolution of all variables in the proposed AdEx model under the influence of a step current *I*. The depicted neuron has the parameters from a Female Group fit neuron.

### 2.3 Peri-PVN Networks Simulations

Both networks have 1000 neurons each, while the peri-PVN has random connections with probability *c_p_* = 0.1, and the CRH^PVN^ network is internally uncoupled. The random connections between sub-networks have probability *c_p_* = 0.1. The peri-PVN network neurons *i* have their dynamics described by the Leaky Integrate and Fire Model (LIF) [29] given by:

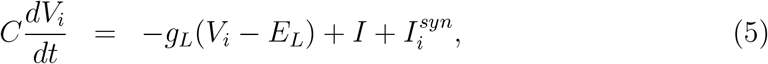

where *C* = 20 pF, *gL* = 0.3 nS, *E_L_* = –70 mV, *I* = 20 pA, the reset voltage *V_r_* = –60 mV, and the peak voltage *V_p_* = –20 mV. The synaptic current 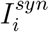 represents the sum of all presynaptic inputs a neuron receives. The synaptic current is given by:

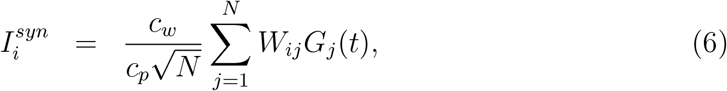

where *c_w_* = 2 is the coupling weight, *N* = 2000 is the total number of neurons in the network, *W_ij_* the adjacency matrix, and *G_j_* is the synaptic current from presynaptic neuron *j*. The function *G_j_* is described by a double exponential function:

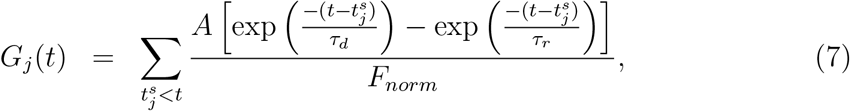

where *τ_d_* = 2 ms is the decay time, *τ_r_* = 1 ms the rise time, *A* =100 pA the amplitude of the synaptic pulse, and 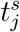 is the *s*th spike fired by the *j*th neuron. The parameter *F_norm_* is a normalization factor given by:

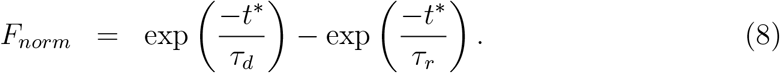

and 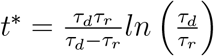. It scales the double-exponential function to have a maximum of 1.

The CRH^PVN^ neurons dynamics are described by Equations (1)–(3), in addition to the synaptic current *I^syn^* to Equation 1. The CRH^PVN^ neurons set of parameters were selected randomly from the Protocol 1 fit models. The external current *I* was set to be equal to their respective rheobase currents *I_rheo_* – 0.1 pA. This was selected to keep the membrane voltage close to the threshold potential resembling a balanced network state [30].

### 2.4 Particle Swarm Optimization - PSO

We used Particle Swarm Optimization (PSO) to search for a set of model parameters that accurately reproduces the CRH^PVN^ neuron current clamp recordings. The PSO algorithm is a gradient-free optimization scheme which is derived from the observations of social interactions in swarming animals and insects. For example, when a swarm of bees searches for a meadow of pollinating flowers, the bees (the particles in PSO) interact autonomously until a bee finds a suitable meadow. This bee then communicates this location to the other bees in the hive [10, 31]. There have been several modifications through the years since the original formulation of PSO by Kennedy and Eberhart [11, 12].

More abstractly, particle swarm optimization describes the movement of a population of agents in a parameter search space. These agents, the particles of PSO, are looking for increasingly better solutions to a specific problem. In particular, the population of agents is seeking to optimize a specific cost function *F_c_*. As the particles move, they exchange information about their own findings, which correspond to more optimal values of *F_c_*. This aids the other particles in their own respective searches. After many iterations and interactions between particles, the population converges towards the position that optimizes the cost function *F_c_*.

The movement equations for each particle *i* at time step *n* are given by [11]:

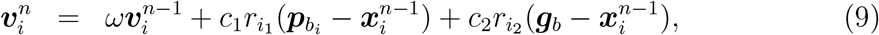

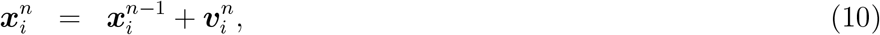

where ***x**_i_* is a *M* dimensional vector of model parameters and ***v**_i_* its velocity. The weight *ω* is commonly referred to as the inertial weight, and is described later. The parameters *c*_1_ and *c*_2_ are positive constants that control the particles’ acceleration. The randomly generated scalars *r*_*i*_1__, and *r*_*i*_2__ are uniform random variables in the range [0, 1], and randomly weigh the two components of the particles movement vector. One component represents a particles own individual search 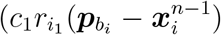 while the other component represents the interaction between members of the swam 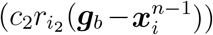. During the search, the particles keep track of their best position, ***p**_b_i__*, which is used to guide their subsequent movements. In addition to this, each particle receives information regarding the best position of the entire swarm, ***g**_b_*, (Figure 1B left).

On the right of Figure 1B, we have schematic illustration of all elements in Equation (9) and the area in grey is the possible new position of the particle (black circle) according to the random variables *r*_*i*_1__ and *r*_*i*_2__. The inertia factor *ω* decreases linearly with time as proposed by [11]:

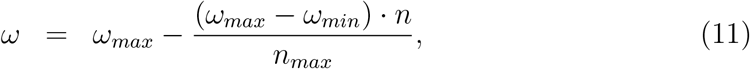

where *ω_max_* = 0.9, *ω_min_* = 0.4 and *n_max_* is the total number of PSO iterations. This modification causes PSO to more heavily favour the independent searches by the particles initially. This increases the swarm’s initial exploration and the chances to find an optimal solution for *F_c_* inside the bounded parameter space by each individual particle. As *ω* decreases, the search strategy switches to exploitation of shared information and the particles communicate more and behave more akin to a swarm. This collective swarm behaviour emerges as the particles search near prior optimal solutions found independently by the separate particles earlier in the optimization.

To avoid instability in the parameter fits, or being confined to the borders of the search space, a maximum and a minimum velocity vector *v_min/max_* are defined. To illustrate the PSO search, Figure 1C shows the swarm movement into the search space of a two-dimensional function *F_c_*. The red cross represents the optimal solution and the red circle the particle with the best position (***g**_b_*). By starting as randomly distributed particles, the swarms position evolves until all particles converge to the solution area. The upper and lower-bounds for all model parameters can be found in Table 1, while all hyper-parameters used for PSO can be found in Table 2.

**Table 1:** Model Neuron Parameters. Lower *L_B_* and upper *U_B_* bounds for all parameters in the PSO search space.

| Model Parameters | $L_B$ | $U_B$ | unit |
| --- | --- | --- | --- |
| $C$ | 1 | 100 | pF |
| $g_L$ | 0.01 | 30 | nS |
| $E_L$ | -80 | -60 | mV |
| $\Delta$ | 0.01 | 30 | mV |
| $a$ | -20 | 20 | nS |
| $V_{T0}^*$ | 5 | 30 | mV |
| $V_{r0}$ | -65 | -45 | mV |
| $V_{p0}$ | 20 | 60 | mV |
| $\tau_w$ | 1 | 2000 | ms |
| $\tau_T$ | 1 | 2000 | ms |
| $\tau_r$ | 1 | 500 | ms |
| $\tau_p$ | 1 | 500 | ms |
| $b$ | 0 | 100 | pA |
| $q$ | 0 | 10 | mV |
| $r$ | 0 | 2 | mV |
| $p$ | -2 | 2 | mV |

**Table 2:** PSO parameters. Hyperparameters of the PSO algorithm.

| PSO Parameters |  |
| --- | --- |
| $P_s$ | 500 |
| $n$ | 500 |
| $c_1$ | 2 |
| $c_2$ | 2 |
| $\lambda$ | 0.7 |
| $v_{min}$ | $-0.2(U_B - L_B)$ |
| $v_{max}$ | $0.2(U_B - L_B)$ |
| $\omega_{min}$ | 0.4 |
| $\omega_{max}$ | 0.9 |

### 2.5 The Cost Function

The PSO process of fitting model neurons involves a complicated, nested, simulation and optimization loop. First, we simulate the voltage trace for each particle in the swam, as distinguished by their set of parameters. This constitutes the inner loop. Each particle in the swarm corresponds to a unique value of the 16-dimensional vector of parameters for the AdEx Model. Then, we calculate the respective cost function *F_c_* by comparison with the CRH^PVN^ data trace, and choose new parameter values for the particles with Equations (9, 10). This constitutes the outer loop.

In our modification of PSO, we use a combination of two error terms to construct the cost function *F_c_*. The first error term, *e*_1_, is associated with the difference between the data traces *V^D^* and model trace *V*. The second error term, *e*_2_, is associated associated with the difference in spiking statistics between the model neuron and the clamped CRH^PVN^ neuron (Figure 1E). These statistics include the total number of spikes (*s*_1_), the amplitude of each spike (*s*_2_), the reset value (*s*_3_), and timing of each spike (*s*_4_).

The first error term, *e*_1_, is given by:

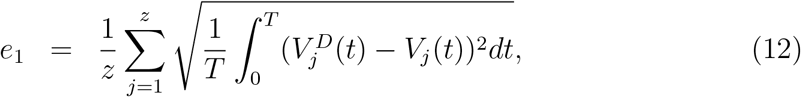

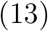

where *z* is the number of recordings for each neuron (at different current clamp levels), *T* the recording time. The equations for *s*_1_, *s*_2_, *s*_3_ and *s*_4_ are given by:

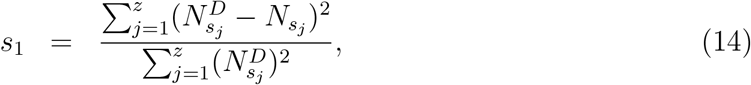

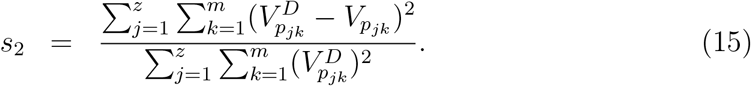

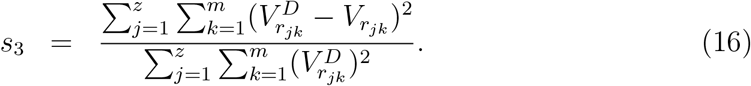

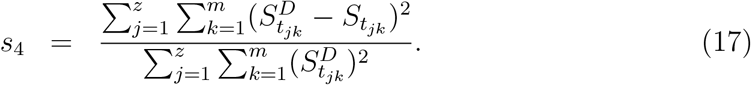

where *N_s_j__* is the number of spikes at the *j*th sweep, *V_p_jk__*, *V_r_jk__*, and *S_t_jk__* are the amplitude, reset value and time of each spike *k* at the *j*th sweep, respectively.

The superscript *D* denotes “Data” in all cases, while model values have no superscript. In Equations (15), (16), and (17) we calculate the error only over the first *m* matching spikes, for example, if data has 5 spikes in a recording *j* and the model presents only 2 then the differences in Equation (15) are calculated only between the first 2 spikes in both. With these components in hand, the function *e*_2_ is defined as follows:

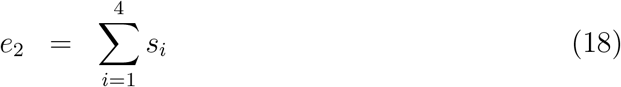

The final cost function is defined as as a linear combination of *e*_1_ and *e*_2_:

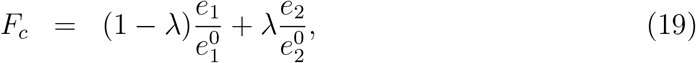

where 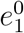 and 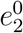 are the errors calculated from the initial random realization of each particle. This normalization is used as an initial reference error and kept fixed through the PSO iterations as a scaling factor for each individual term. The constant λ can vary in the interval [0, 1], thereby balancing the cost function terms. The value of *λ* = 0.7 was fixed for all fits, and this was determined by an initial simulation analyzing the impact of lambda in *F_c_* (Figure 3).

**Figure 3:**
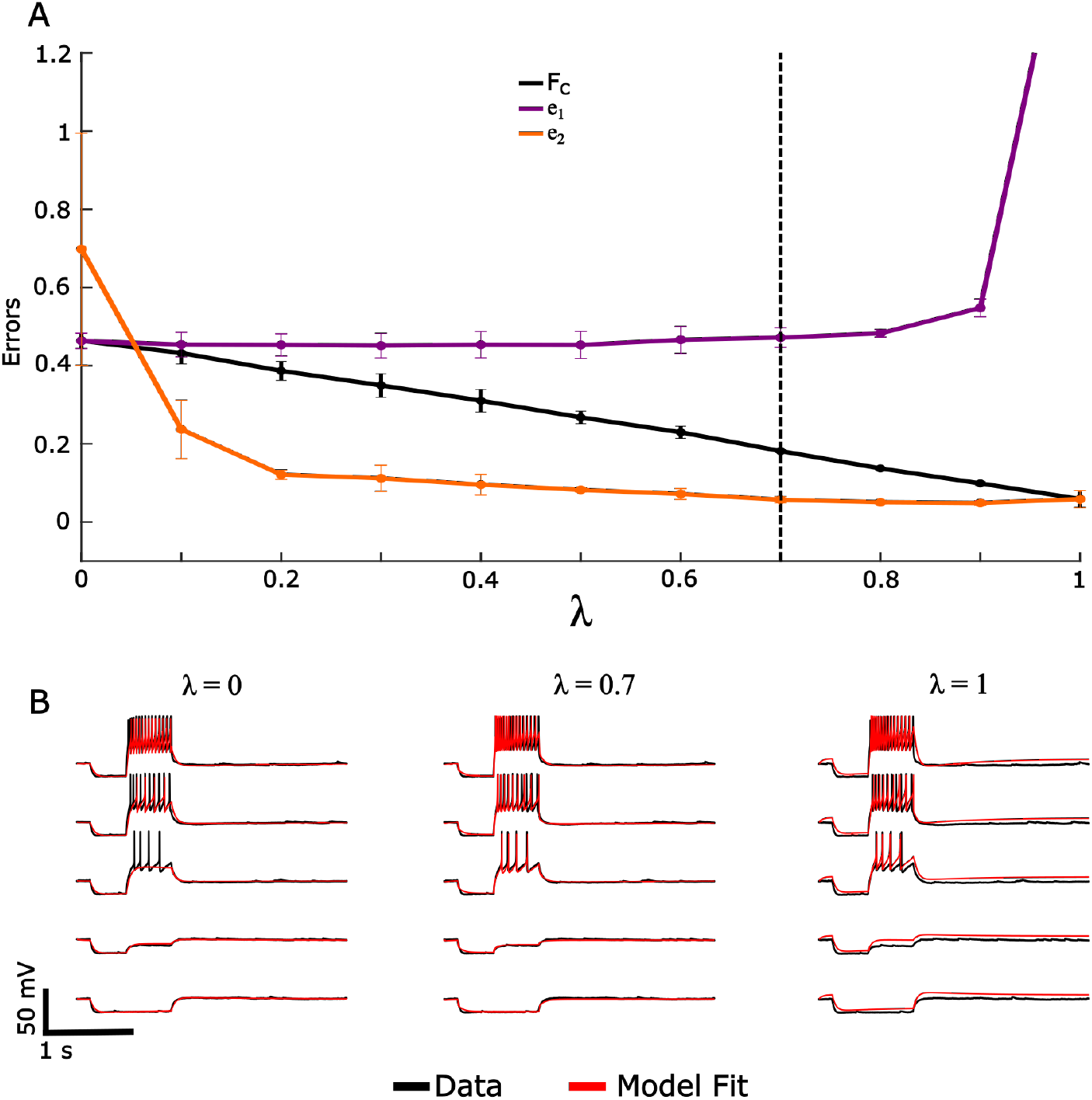
Single neuron fit cost function as a function of λ. **(A)** The values of *F_c_* as *λ* increases (black line), the associated *e*_1_(purple) and *e*_2_ (orange) errors. The dashed line depicts the value used for our PSO fits (λ = 0.7). **(B)** The fits for three different λ values, for λ = 0 the spikes are not well fit and for λ =1 the spike features are well reproduced but not the subthreshold dynamics. The chosen value λ = 0.7, which used globally for all neurons, presented a good balance between spike features and subthreshold membrane potential. The analysis was made with the model data of a fit neuron from the Female Group sample, and for each λ in the interval [0, 1] the average cost function and errors were calculated over 5 trials.

Example fits (Figure 1D) to a single current clamp recording and the associated errors (Figure 1F) *e*_1_ and *e*_2_ show a general decrease in the error for increasing the amount of particles, and the amount of iterations of PSO. While increasing the amount of swarm particles and iterations improves fitting, this also increases the simulation time and computational cost.

To prevent biologically unrealistic values, the model parameters were fit with lower *L_B_* and upper *U_B_* bound constraints (Table 1). Further, we added the constraint 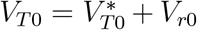 to ensure that the resting threshold voltage will always be above the reset voltage, thereby avoiding pathological bursting/spiking. The PSO parameters can be found in Table 2. For each neuron, 32 repetitions (runs) of the PSO fitting were performed, all of them normalized in relation to the same initial random realization (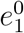 and 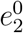) to enable comparability. In all runs, the particles’ initial positions ***x***_0_ were randomly distributed inside the limits of the parameters space, and their initial velocity was fixed to ***v***_0_ = 0.1***x***_0_.

For each neuron we separated some current clamp recordings for the fitting or training (red traces on Fig 1H) and left some for testing the solution (blue traces on Fig 1H). Critically, the testing set is not used to compute the cost function, *F_c_* during the PSO and is left out as an independent comparison of model quality. In Protocol 1, neurons were separated into five current clamp values for PSO training and two for testing. For Protocol 2, we used nine clamp values for training and three for testing per neuron. It is important to highlight that the PSO algorithm, as implemented here, searches for a set of parameters that fit all the training recordings at once and not a different parameter set for each recording. This allows the models to generalize to the test sets.

Any model (that is not overfit) will have some error associated with it, as such, the criterion that *F_c_* = 0 is not a suitable stopping measure for PSO. Thus, the total number of iterations *n* becomes the criteria for stopping PSO and comparing *F_c_* over particles. Once stopped, we use the global minimum ***g**_b_* as the parameter value for the CRH^PVN^ model.

Figure 1G shows the pair of errors *e*_1_ and *e*_2_ for all fit neurons of Female Group Housed sample and the colour scale shows the *F_c_* associated value. The *F_c_* value may vary between neurons since each fit is made independently, therefore even with different *F_c_*, the models can accurately mimic the experimental data (Fig 1H).

### 2.6 Dynamical Features and Bifurcation Analysis

Existing bifurcation analyses of the AdEx model neurons considered here can provide detailed information about properties not directly captured from the electrophysiological data. Despite the modifications we have made to the AdEx model, the work of [32] in deriving the analytical equations for the fixed points, and bifurcations of the AdEx model is directly applicable. The reason for this applicability is that all of the modifications we have made to the AdEx model are spike-dependent, and thus impact the non-smooth spiking/Fillipov dynamics [32], rather than smooth dynamics near the equilibrium.

The AdEx model can have up to two equilibria, which are stable and unstable. These are given by:

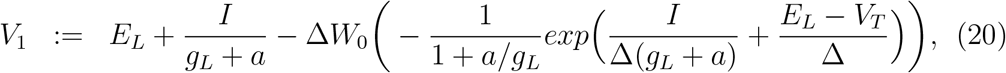

and

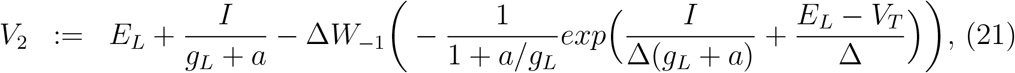

where *W*_0_ and *W*^-1^ are the principal and real branch of Lambert *W*(*x*) function respectively [32]. This function is defined as the solution to the equation:

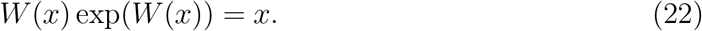

When both of these equilibria exist, *V*_1_ is stable while *V*_2_ is unstable [32].

The excitability type of a neuron defines its responses to a variety of external currents *I*(*t*), and the neurons’ transition to spiking. This transition to spiking occurs formally through a bifurcation [32]. For 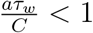, the neuron undergoes a saddle-node bifurcation where the equilibria *V*_1_ and *V*_2_ merge and eliminate each other. This is classified classically (by Hodgkin and Huxley) as Type-1 excitability [33], where arbitrarily low firing rates are possible. For *aτ_w_*/*C* > 1, the neuron has Type-2 excitability and undergoes a subcritical Andronov-Hopf bifurcation where the fixed point *V*_1_ becomes unstable by colliding with an unstable limit cycle. As a result, Type-2 neurons have a discontinuous firing-rate as a function of their input current. Type-2 Neurons are not able to produce arbitrarily low firing rates. Further, saddle node and Hopf bifurcations display different neural sub-threshold dynamics. In the former, signals are integrated by the membrane, while in the latter, the membrane has a characteristic resonance frequency in which it maximally responds. When integrators receive a (small) current, there is a membrane voltage increase followed by an exponential decay to the resting state. Resonators, on the other hand, decay with a damped frequency.

The rheobase current *I_rheo_* is classically defined as the minimum current of infinite duration necessary to trigger a spike [34]. As the current intensity increases, the rheobase corresponds to the value where *V*_1_ loses stability. For integrator neurons, this is given analytically by the saddle-node bifurcation point [32]:

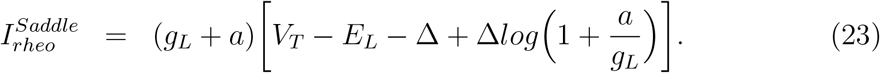

The rheobase of resonator neurons, on the other hand, corresponds to the Hopf bifurcation point. This is given by:

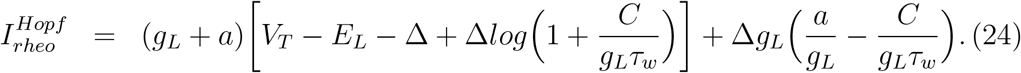

Those quantities can also be calculated numerically via direct simulation as done here, or with numerical bifurcation software such as MATCONT/XPPaut [35], [36]. The direct simulation values show excellent agreement with the analytical equations from [32], (Figure 5).

**Figure 4:**
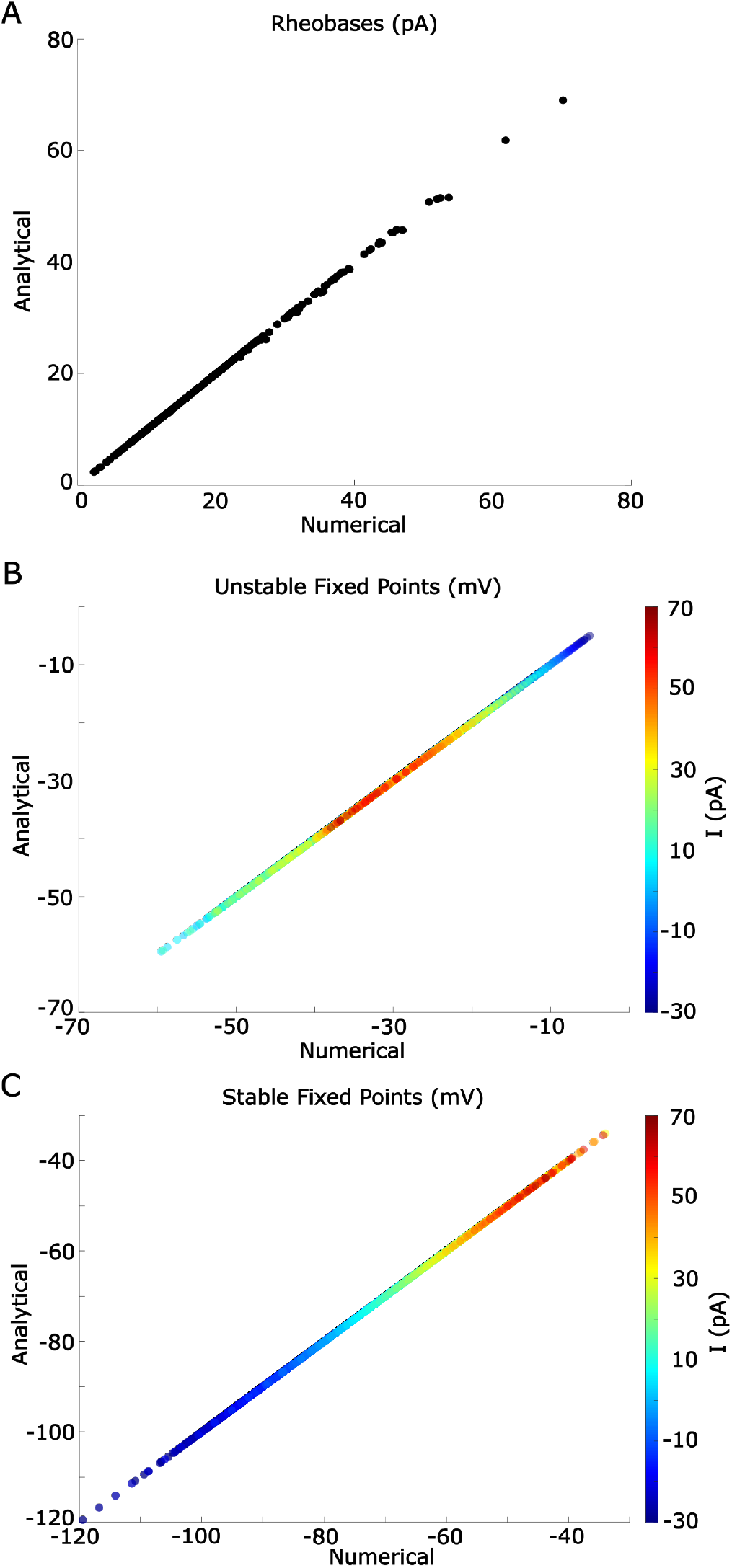
Numerical and analytical dynamical features. Comparison of the dynamical features obtained numerically from the model and the analytical solutions from [32]. **(A)** The rheobase currents for all neurons in Protocol 1. **(B)** Unstable equilibrium for all neurons from protocol 1 (as in A). **(C)** Unstable equilibrium, as in (B). The equilibria is measured over the [-30,70] pA range.

**Figure 5:**
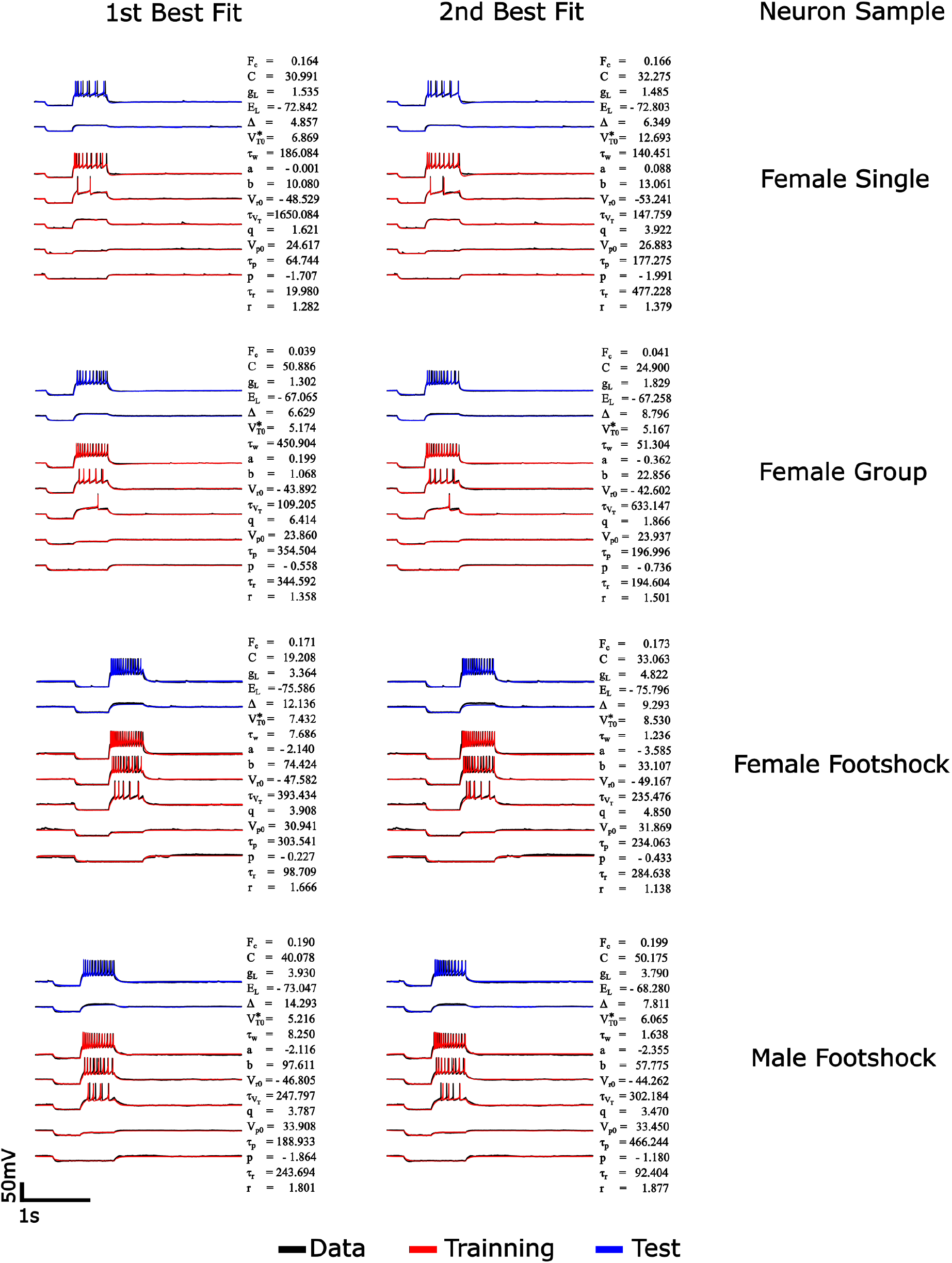
First and second best fits for a subset of neurons. Example fits of a neuron from each sample in Protocol 1. On the left we show the neuron first best fit and on the right the second best fit, according to their respective *F_c_*, together with the parameter set. For units of variables, see Table 1.

## 3 Results

### 3.1 Particle Swarm Optimization can Accurately and Rapidly Fit CRH^PVN^ Neurons

To date, the (CRH^PVN^) of the Paraventricular (PVN) nucleus have yet to be analyzed and modeled in a data driven way. To investigate their dynamical features across mouse sex and stress condition, we fit all 248 patched neurons (Figure 1A) to a modified version of the Adaptive Exponential Integrate-and-Fire neuron model:

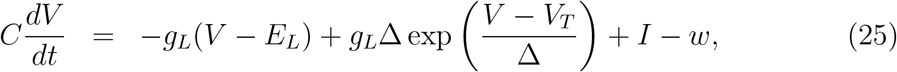

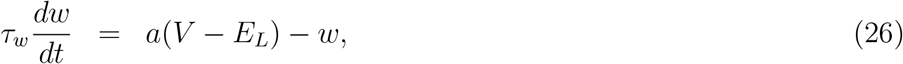

where *V* is the neurons voltage, *w* acts as an adaptation variable. The modifications made include an adaptive threshold, an adaptive voltage peak (*V_p_*), and an adaptive voltage reset (*V_r_*). The voltage peak, reset, and threshold change with regards to the spiking history (Section 2.2), which was a dynamic observed in the patched CRH-neurons.

In total, the modified AdEx model consists of 16 parameters, which were fit with particle swarm optimization (PSO). The parameters in PSO correspond to particles which move collectively to minimize a cost function, *F_c_*, which dictates the quality of a fitted model to a patched neuron (Section 2.5). Each patched neuron’s recorded voltage traces were separated into training and test traces. The training traces were used in the PSO optimization to minimize *F_c_*, while the test traces were held out to determine the quality of fit (Section 2.1). As the particles find better fits, they communicate that information amongst each other in a collaborative effort to optimize *F_c_* (Figure 1B-C). As more particles are used, or more iterations of PSO are applied, the algorithm converges towards models that fit the CRH^PVN^-neuron’s voltage with low values of the cost function *F_c_* (1D-F).

The cost function itself consists of a term which penalizes the squared difference between the model and a patched neuron’s voltage trace, and a term which penalizes the timing, number, and characteristics of spikes (Section 2.5, Figure 1E). All neurons were successfully fit, albeit with different values of these two error components in *F_c_* (Figure 1G). However, even the worst fit neuron (according to the *F_c_*) shows a remarkable accuracy in reproducing a patched CRH^PVN^ neuron’s subthreshold and spiking characteristics for both training traces and test traces. Collectively, these results show that PSO, paired with the right cost function, is a viable tool for fitting to a neuron’s voltage traces and spike times recorded with routine current clamp experiments.

### 3.2 Particle Swarm Optimization Leads to Multiple Solutions for Any Given Neuron

With the ability of PSO to accurately fit AdEx models to CRH^PVN^ neurons firmly established, we next wondered if these fits were in anyway unique. This was motivated by prior results where multiple solutions or sets of model parameters for other neurons were found [15–17].

We observed that multiple sets of parameters can reliably reproduce CRH^PVN^-voltage traces. The individual parameters themselves often differed radically from the 1st to 2nd best fits (Figure 5). To study the non-uniqueness of the PSO generated models, we selected for each CRH^PVN^ neuron the two best fits over the 32 PSO runs according to the cost function *F_c_*. While the raw parameters differed substantially, Principal Component Analyses (PCA) of parameters yield similar low dimensional projections (Figure 6A), and the correlations between the individual parameters were largely preserved from the first to second best fits (Figure 6B). This indicates that the relationship between parameters was preserved as some type of high dimensional parameter manifold, but not necessarily the parameters themselves. Further, both the first and second best fits are significantly correlated regarding their first principal component (Figure 7).

**Figure 6:**
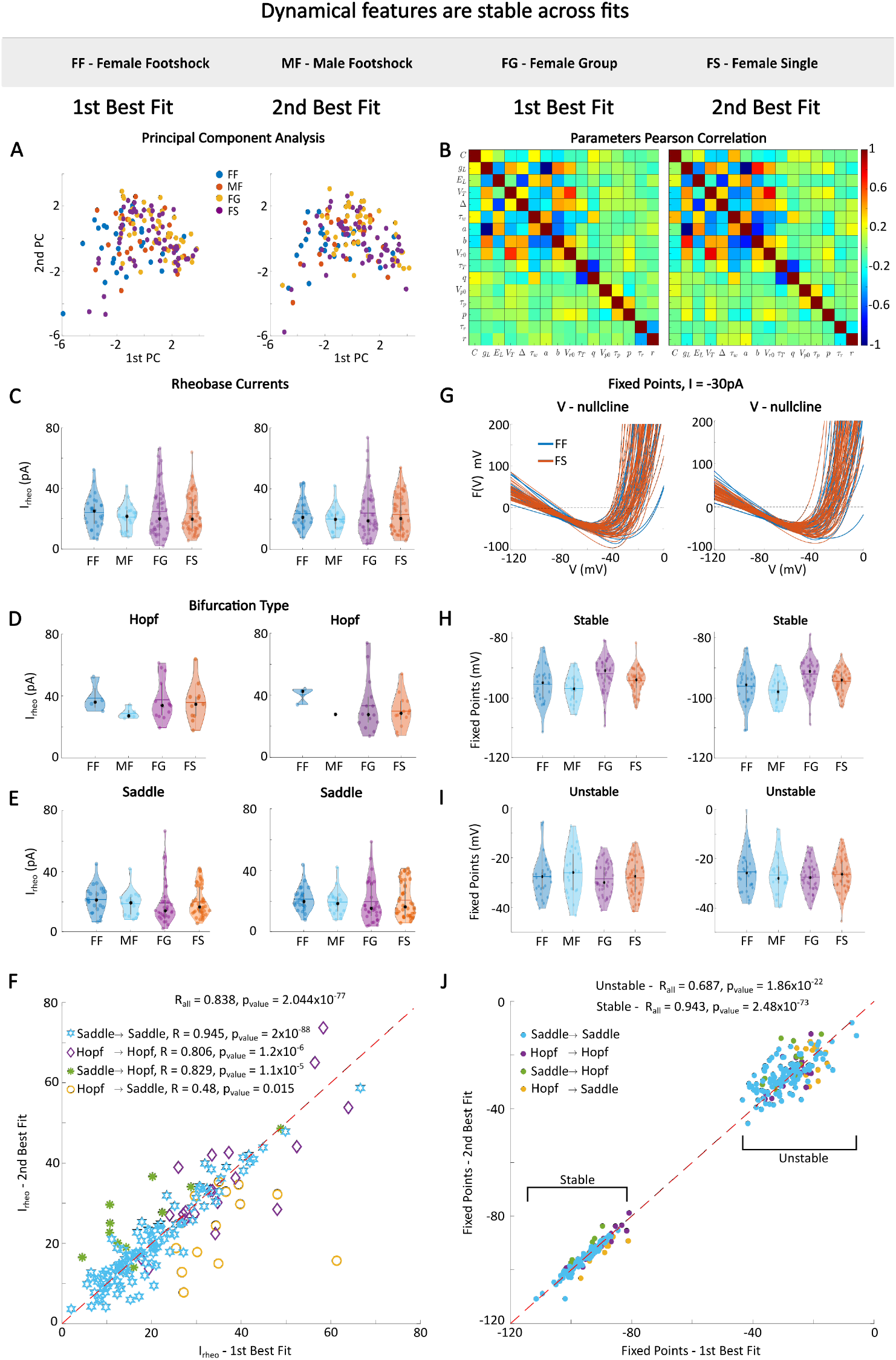
Comparison between the 1st and 2nd best fits parameters and dynamical features. **(A)** PCA of model parameters from all neurons in Protocol 1. **(B)** Pearson correlation between model parameters into the 1st (left) and 2nd (right) best fit of all sample neurons. **(C)** Violin plots with the distribution of rheobase currents per sample, the black dots are the median and the horizontal lines the mean. **(D)** Rheobase distribution for Hopf neurons for 1st (left) and 2nd (right) best fits. **(E)** Rheobase distribution for saddle-node neurons for 1st (left) and 2nd (right) best fits. **(F)** Pairs of rheobase currents of the 1st and 2nd best fit. Symbols denote the type of bifurcation and if the bifurcation changed between 1st and 2nd best fits. **(G)** V-Nullcline of all neurons in Female Footshock and Female Single samples for *I* = −30 pA, the intersection points of the red/blue lines with the dashed line (*F*(*V*) = 0) provides the fixed points values numerically. **(H)** Distribution of stable equilibria for 1st (left) and 2nd (right) best fits. **(I)** Distribution of unstable equilibria for 1st (left) and 2nd (right) best fits. **(J)** the same as in (F) with respect to the fixed points.

**Figure 7:**
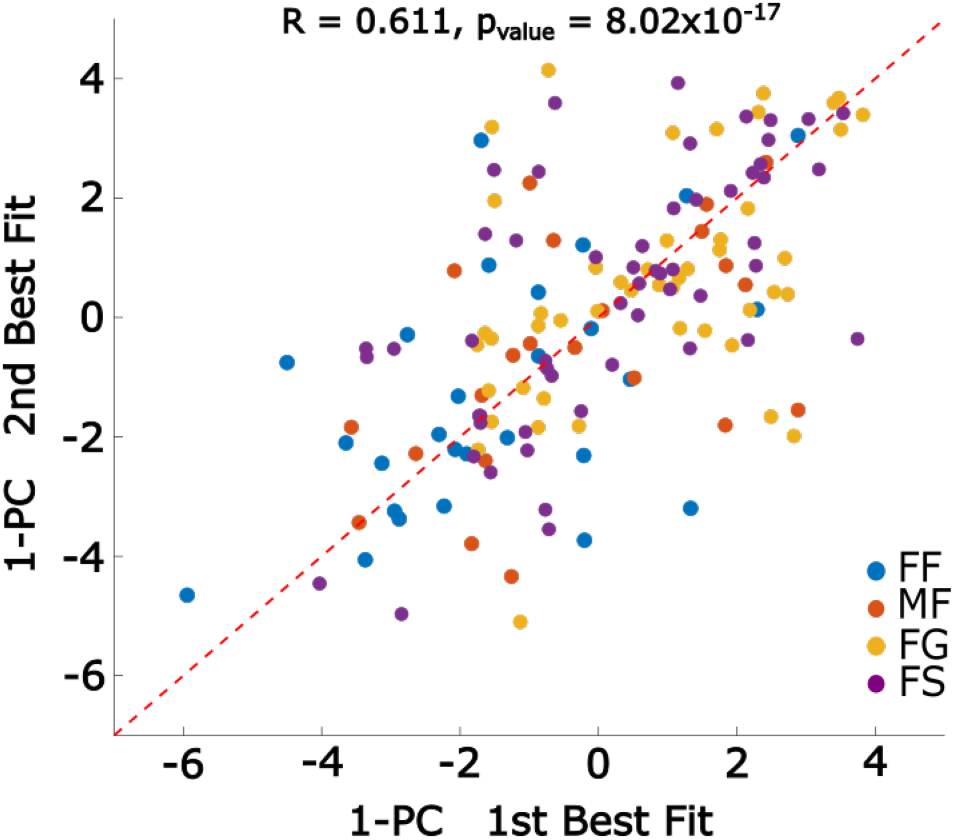
First principal component analysis of all fit models’ parameters. Correlation between the 1-PC of parameters from the first and second best fits of all neurons in Protocol 1. The colors represent each sample in Protocol 1.

With the parameters themselves inconsistent between the 1st, and 2nd best fits, but the correlation between parameters preserved, we next wondered if the dynamical features of the isolated CRH^PVN^ neurons were preserved from the 1st to the 2nd best model fits. Thus, we considered the following dynamical features: the rheobase current, the bifurcation type, and the equilibrium points. We found that the distributions for all of these dynamical features were stable (Figure 6C-J) from the 1st to the 2nd best fits. Further, the explicit values themselves were highly correlated from 1st to 2nd best fits (Pearson’s *R* of 0.84, *p* = 2.02 × 10^-77^ for rheobases, ***R*** = 0.69 and *p* = 1.86 × 10^-22^ for the unstable fixed points, and *R* = 0.94, *p* = 2.48 × 10^-73^ for the stable fixed points. Finally, only a minority of points were found to transition between bifurcation types from the 1st to 2nd best fits, likely representing models that could be fit better with increasing iterations of PSO.

Our results show that PSO provides multiple sets of parameters that can reliably reproduce the behaviours of CRH^PVN^ neurons. Despite these non-unique solutions, the dynamical features are largely stable across fits.

### 3.3 Particle Swarm Optimization is Sensitive to Different Current Clamp Protocols

The CRH^PVN^ neurons were recorded under two different current clamp protocols. In the first protocol, the neurons received an initial hyperpolarizing current to current immediately followed by a depolarizing current of increasing amplitude. This was done to ensure voltage-gated ion channels were not in the inactivated state, and able to open in response to depolarization (Figure 8A left). In the second protocol, neurons received a depolarizing current without an initial hyperpolarizing prepulse (Figure 8A right).

**Figure 8:**
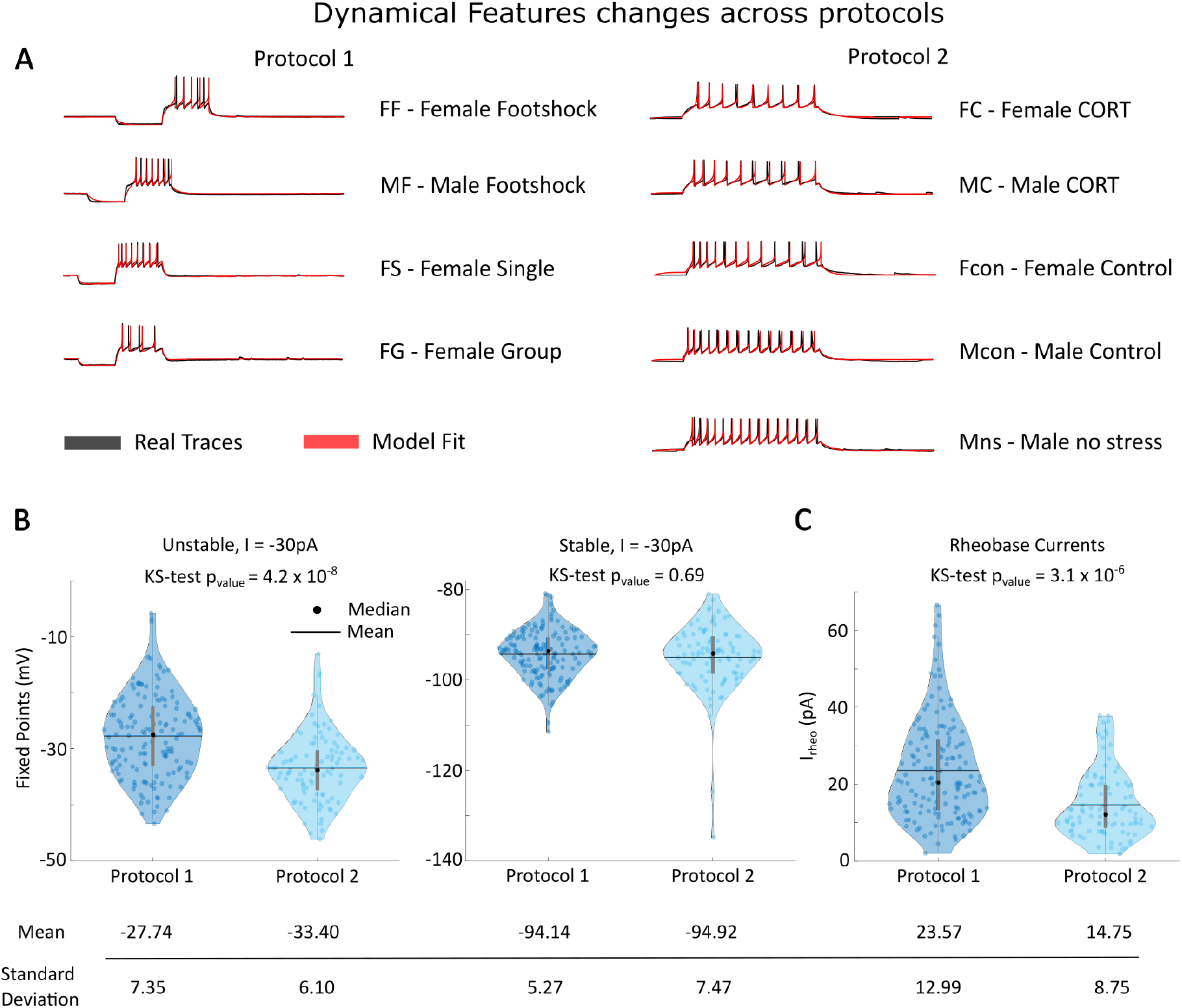
Differences in cell dynamics due to protocols. **(A)** Protocol 1 and 2 traces characteristics, in black the data traces and in red the fit models. **(B)** Distribution of equilibria for all neurons in both protocols. On the left we have the unstable fixed points and on the right the stable ones, both inferred for an external input *I* = −30 pA. **(C)** Rheobase currents distributions across protocols. On the botton of (B) and (C) are depicted the mean and standard deviation of all the respective quantities.

The PSO search was able to perform a reliable reconstruction of both protocols’ recordings, with largely similar qualities of fits (Figure 8A). However, the protocols themselves lead to different dynamical features. In particular, the unstable equilibrium is substantially and significantly lower for the second protocol (KS-test *p* = 4.4 × 10^-8^, Figure 8B left). In contrast, the stable fixed points have similar distributions for both protocols (KS-test, *p* = 0.69, Figure 8B right). The rheobase currents was also found to have a lower value on average (KS-test *p* =3.1 × 10^-6^, Figure 8C). These results shows that the dynamical features are not stable across protocols, and also suggest that different current clamp protocols can influence or change neuronal properties. The PSO algorithm was able to fit neurons, however the fits are sensitive to the specifics of the current clamp procedure used, which may mask or unmask different elements of the neuronal dynamics.

### 3.4 Bifurcation Analysis of CRH^PVN^ Model Neurons

The type of bifurcation defines how a neuron responds to a current ramp, as well as how the membrane potential returns to the resting state after receiving an excitatory or inhibitory post synaptic potential. Thus far, PSO has revealed two distinct sub-populations of CRH^PVN^ neurons: CRH^PVN^-resonators and CRH^PVN^-integrators, which are distinguished by subcritical-Hopf and saddle-node bifurcations, respectively. The proportions of Hopf and saddle neurons in each sample can be observed on Figure 9A. The Protocol 1 samples contain approximately 23% CRH^PVN^-resonators, and Protocol 2 samples contain 16% CRH^PVN^-resonators.

**Figure 9:**
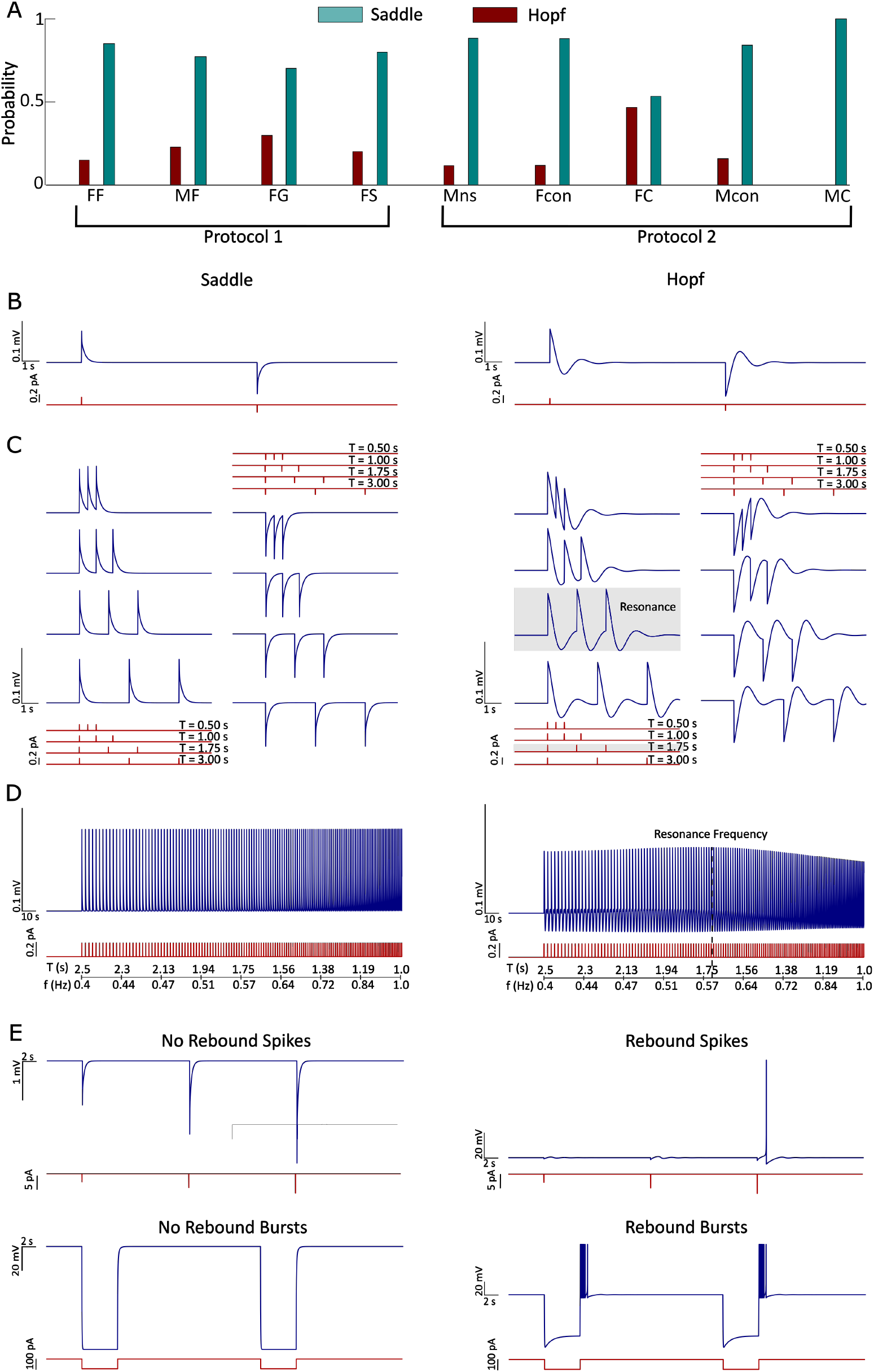
Dynamical features of CRH-Resonators and Integrators. **(A)** Proportion of saddle (CRH^PVN^-integrators) and Hopf (CRH^PVN^-resonator) neurons in each sample. **(B)** Saddle (left) and Hopf (right) neurons sub-threshold response to a excitatory and inhibitory current pulse. **(C)** Effect of excitatory and inhibitory train of pulses with different periods *T* = [0.5 s, 1.0 s, 1.75 s, 3.0 s]. Each neuron received a baseline current *I_b_* = *I_rheo_* ± 0.4 pA, and the external current pulses have 0.2 pA magnitude. **(D)** Sub-threshold dynamics response to a increasing frequency ([0.4 — 1] Hz) current pulse. The CRH^PVN^-resonator neuron here present resonance around 0.6 Hz. The CRH^PVN^-resonator received a baseline current *I_b_* = *I_rheo_* — 0.25 pA. **(E)** Neural response to a high magnitude (100 pA) pulse (top) and 5 s step (botton) currents. Rebound spikes/bursts can be observed in Hopf neurons exclusively. A baseline current of *I_b_* = *I_rheo_* – 0.1 pA was used. For all simulation in (B-E) we used two selected model neurons from FF sample.

With these neuronal parameters in hand, we wondered how CRH^PVN^-integrators and CRH^PVN^-resonators would respond to currents that were outside of the current clamp protocols considered experimentally. CRH^PVN^-integrators display a monotonic decay to the resting membrane potential when receiving an excitatory or inhibitory pulse of current (Figure 9B left). As all integrators do, the CRH^PVN^-integrator population linearly sums up the excitatory or inhibitory currents, followed by a monotonic decay (Figure 9C left). The faster a sequence of pulses arrives at a synapse, the greater the deviation of the current (Figure 9D left) with a monotonic increase in the voltage deviation when these pulses are excitatory. CRH^PVN^-integrators also respond with strict hyperpolarization for phasic and tonic inhibition (Figure 9E).

CRH^PVN^-resonators however display prominent subthreshold oscillations when receiving excitatory or inhibitory pulses of current (Figure 9B right). These pulses display a prominent resonance phenomenon (Figure 9C-D right) at ultra-slow frequencies (comparizon of numeric and analytical resonance frequencies and distribution depicted in Figure 10 [18, 19]) with a mean frequency of 0.45 Hz ± 0.25 Hz. Further, these neurons also display prominent rebound spikes with sufficiently strong hyperpolarizing phasic or tonic currents (figure 9E right).

**Figure 10:**
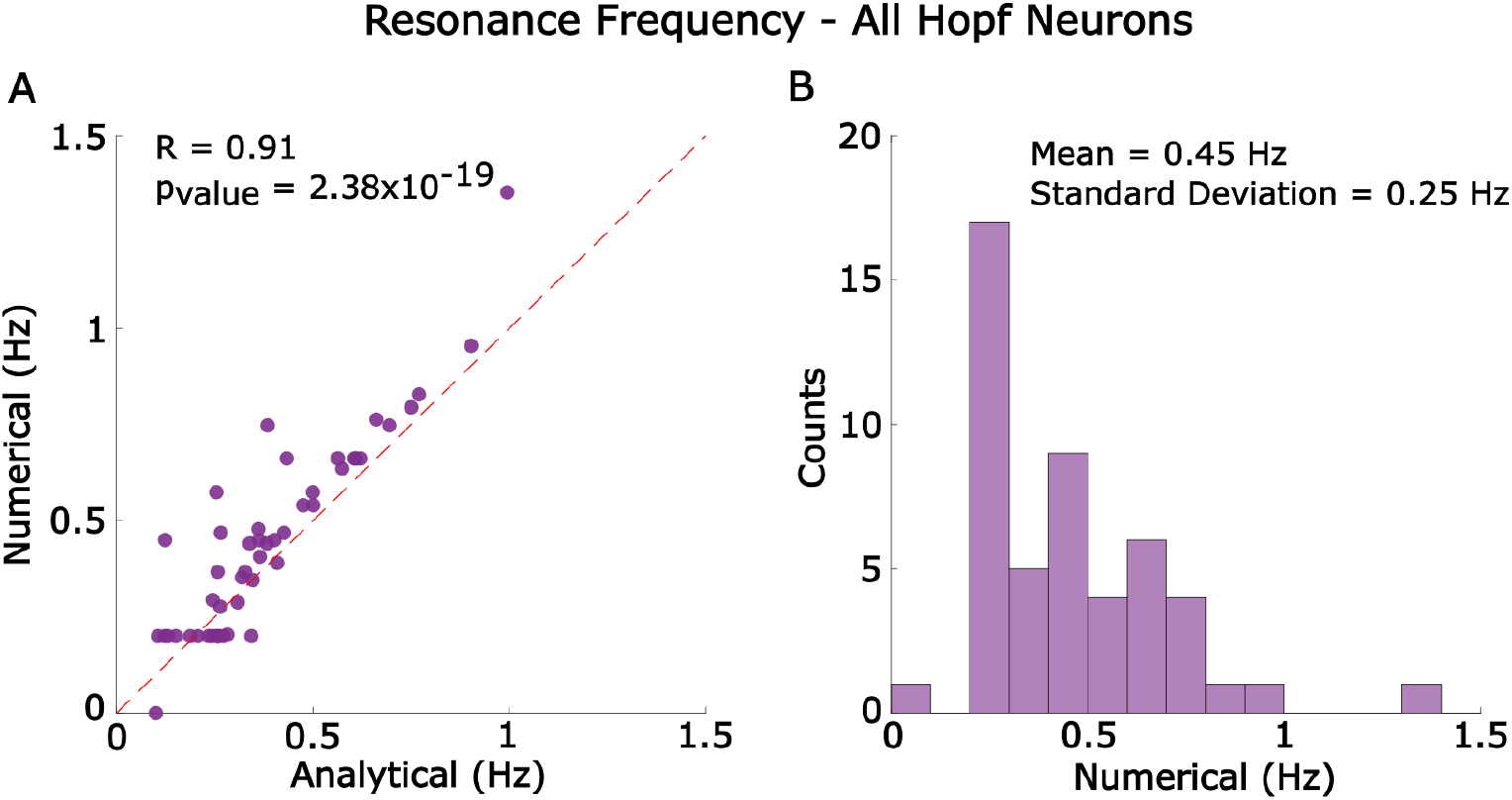
Numerical and analytical resonance frequencies. **(A)** Comparison of the resonance frequencies obtained analytically and numerically, and **(B)** Distribution of numerical resonance frequencies. To find the numerical results all Hopf neuron received a baseline current *I_baseiine_i__* = *I_rheo_i__*, — 0.25*pA* and a pulse current I = 0.2pA with an linearly increasing frequency in the range [0.1, 2] Hz in a 50s simulation time, the resonance frequency was selected from the voltage peak with the highest amplitude. The analytical resonance frequency was obtained from imaginary part of the eigenvalue at the bifurcation point [18, 19].

### 3.5 The CRH^PVN^-Resonators Can Maintain Baseline PVN Firing Even with Blanket Inhibition

The heterogeneity of parameters and the behavioral richness of the CRH^PVN^ neuron lead us to investigate how networks of CRH^PVN^-integrators and resonators would respond to a mixture of excitatory and inhibitory inputs. We simulated networks of 500 uncoupled neurons, either entirely CRH^PVN^-integrators or CRH^PVN^-resonators, receiving independent randomly generated currents *I_i_*(*t*) (Fig 11A). These neurons were generated by randomly selecting from models fit with Protocol 1 from the resonator or integrator class. The current *I_i_* was generated from a Poisson distribution with a mean frequency of random pulses. These pulses were selected to be excitatory or inhibitory according to a specified probability (Figure 11B). We studied and compared the Peristimulis time histogram (PSTH) mean activity of CRH^PVN^-integrator and CRH^PVN^-resonator networks for increasing input frequencies (Figure 11C). We varied the input frequency of these pulses for both the integrator and resonator networks. The CRH^PVN^-resonator network had higher frequencies of firing for all input frequencies tested (1 Hz, 10 Hz, and 50Hz), and for all excitation to inhibition ratios (Figure 11D). The CRH^PVN^-resonator networks also had the ability to sustain firing activity even under total inhibitory inputs. This non-zero firing rate under total-inhibition in CRH^PVN^-resonators is the result of the post-inhibitory rebound displayed by these neurons. These results suggest that a population of CRH^PVN^ neurons with a mix of integrators and resonators could sustain a minimum spike rate, and therefore possibly even a minimum release of CRH^PVN^, even under overwhelming inhibition.

**Figure 11:**
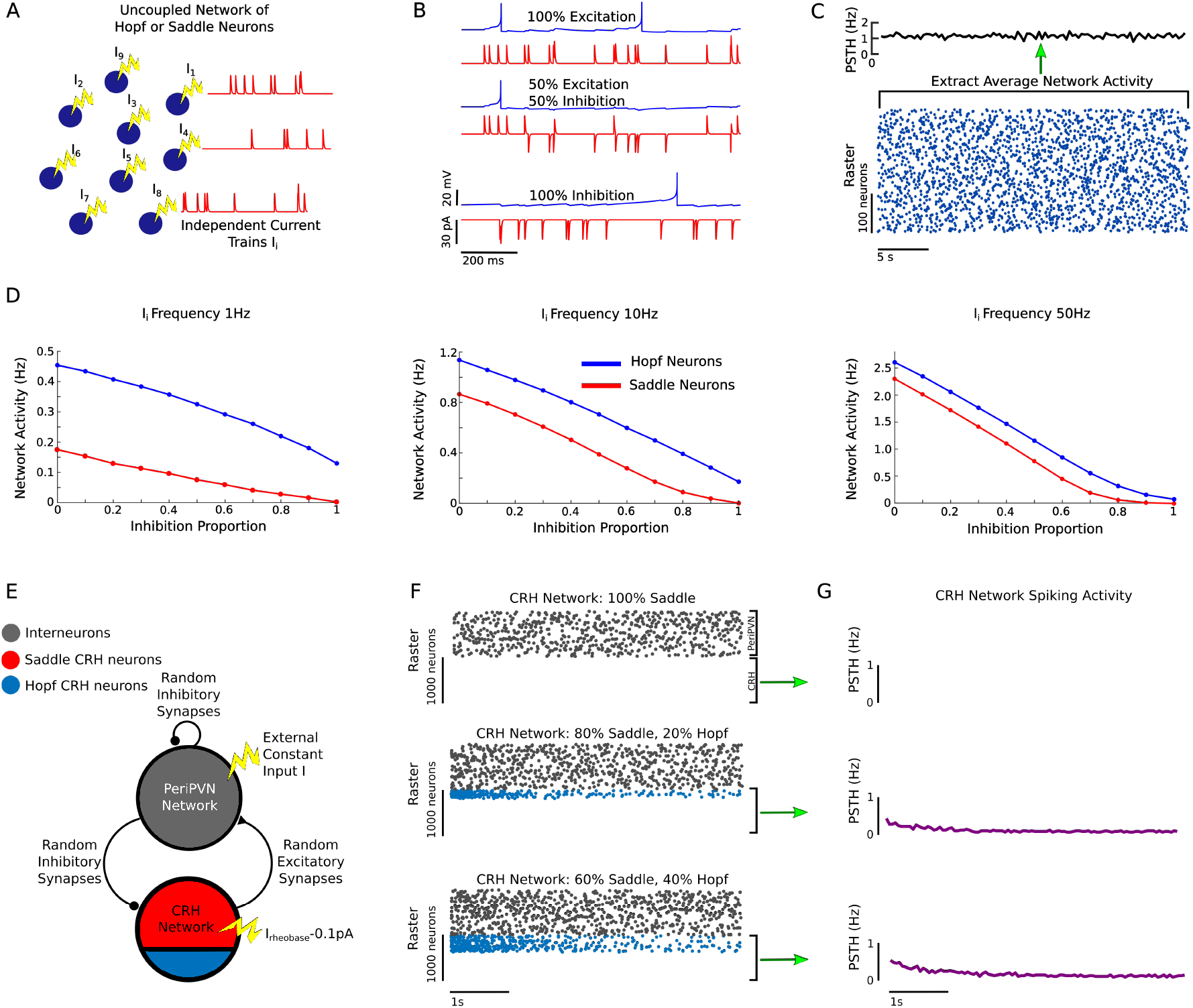
Network dynamics of CRH^PVN^ neurons. **(A)** Schematic illustration of uncoupled neurons in the network, each one receiving an randomly generated and independent input current. **(B)** Response of a neuron to the impulse signal for different proportion of excitation and inhibition. **(C)** Raster and Peri-Stimulus-Time-Histogram (PSTHJ) of a 500 CRH^PVN^-resonator network. In this example, the external current has a proportion of 50% excitation and 50% inhibition. **(D)** Mean activity of CRH^PVN^-resonator (blue) and CRH^PVN^ integrator (red) networks as the proportion of inhibition increases for average current frequencies of 1 Hz (left), 10 Hz (middle) and 50 Hz (right), the magnitude of the pulses were fixed to |*I*| = 30 pA in all simulations. **(E)** Schematic illustration of the biologically based peri-PVN/CRH^PVN^ network. The peri-PVN network is composed of 1000 LIF interneurons in a balanced inhibitory regime with 1000 excitatory uncoupled neurons randomly selected from the fit PSO models (Protocol 1), the proportion CRH^PVN^-integrator (red) and CRH^PVN^-resonator (blue) neurons may vary according to the simulation. The connections between both networks are random (0.1 probability). Each peri-PVN neuron receives a constant current *I* = 20 pA and the CRH neurons receive each their respective rheobase current decreased by 0.1 pA, resembling a balanced state. **(F)** Raster plots for different proportion of resonator neurons in the CRH network (top) 100% integrators, (middle) 80% integrators /20% resonators, and bottom 60% integrators/40% resonators. **G** PSTH of the CRH network for the conditions described in (F).

To confirm this result in a more biologically plausible scenario, we studied the activity of a network formed by two coupled sub-networks: a peri-PVN inhibitory network of interneurons, and a population of CRH^PVN^ excitatory neurons network consisting of CRH^PVN^-integrators and resonators (Figure 11E). We considered three network configurations (Figure 11F): 100% CRH^PVN^-integrators, middle 80% integrators /20% resonators, and 60% integrators /40% resonators. Once again, the peri-PVN coupled CRH^PVN^ network only displays activity through the CRH^PVN^-resonators at rest via a rebound spiking (Figure 11F). These neurons also have a strong initial transient response which stabilizes over long time-scales to superthreshold firing (Figure 11G). Collectively, these network simulations suggest that a small population of CRH^PVN^-resonators may be critical for maintaining baseline firing in the HPA-axis.

## 4 Discussion

The mammalian stress response is critically controlled by CRH^PVN^ neurons in the paraventricular nucleus (PVN) of the hypothalamus. We found that these neurons could be rapidly and accurately modeled with a modified Adaptive-Exponential Integrate- and-Fire neuron model where the parameters were fit with particle swarm optimization. The parameters found by PSO were non-unique for each individual neuron, but the dynamical features, which are non-linear functions of the individual parameters were unique and highly correlated across the 1st and 2nd best fits. The CRH^PVN^-neuron models could be classified as two distinct classes: CRH^PVN^-resonators, which undergo a Hopf bifurcation and display a slow (≈ 0.5*Hz*) resonance frequency, and CRH^PVN^-integrators, which undergo a saddle-node bifurcation. Network simulations showed that CRH^PVN^-resonators were critical for maintaining the baseline firing of the CRH^PVN^ population in realistic models of CRH^PVN^ networks with peri-PVN inhibitory inputs.

The PSO fits determined non-unique solutions to the parameters required to fit a CRH^PVN^ neuron. This, however, is not a consequence of using PSO itself, but rather seems to be a general feature of neuron modelling, and potentially neurons themselves. Indeed, prior analyses of the somatogastric ganglion of crustaceans shows multiple synaptic, and even network parameters in a neuron model can lead to nearly identical dynamical behaviours [15–17]. Our results here show the mammalian equivalent in the stress circuit and lead directly to the hypothesis that animals potentially exploit multiple parameter solutions in single neurons to maintain homeostasis in the HPA axis.

Our results also show that CRH^PVN^-neurons have two different subtypes: CRH^PVN^-resonators with ultra-slow resonance frequencies, and CRH^PVN^-integrators. The CRH^PVN^ resonators maximally responds to very slow frequencies on average, but with a diversity of resonance frequencies over approximately an order of magnitude. A dedicated sub-population of these CRH^PVN^-resonators could potentially act as a sequence of independent clocks, calculating and responding to periodic inputs at behavioural time scales. Interestingly, slow-resonance has been documented in non-orexin cells in the lateral hypothalamus [20], suggesting that this may be a more general mode that can selectively and differentially recruit different hypothalamic subsets via oscillations of specific frequencies.

CRH^PVN^ neurons display a diversity of behaviors that can be achieved with non-unique electrophysiological parameters per neuron. Future work may directly confirm the existence of slow-resonance with targeted patching with oscillatory input currents, and potentially elucidate the role of slow-resonance frequencies under in vivo conditions.

## Aditional Information

### Data availability statement

The data that support the findings of this study are available from Jaideep Bains laboratory upon request.

### Author contributions

W.N and J.B conceived and provided resources for the study. N.P.R, D.V and S.L performed the experiments. E.L.L and W.N analysed the data, developed the simulations and wrote the manuscript. All authors approved the final version of this manuscript and agree to be accountable for all aspects of the work in ensuring that questions related to the accuracy or integrity of any part of the work are appropriately investigated and resolved. All persons designated as authors qualify for authorship, and all those who qualify for authorship are listed.

### Funding

The projects are supported by an NSERC Discovery Grant and a CIHR FDN award, and a Hotchkiss Brain Institute start-up fund, N.P.R. is supported, in part by the Cumming School of Medicine’s “Leaders in Medicine” program.

